# Bacteriophage to Combat Biofilms in Hospital Drains

**DOI:** 10.1101/522227

**Authors:** Jenny Yijian Huang

## Abstract

**Background:** According to the World Health Organization, nearly 15% of all hospitalized patients worldwide acquire nosocomial infections. A particular area of concern for bacterial build up in hospitals is sink drains. The moist, microbiologically active environment of drains promotes the formation of biofilms that are difficult to target with standard chemical disinfectants. Bacteriophages, however, show potential to be used as disinfecting agents in hospital drains. Not only do bacteriophages increase in titer as they infect, spreading to hard-to-reach surfaces, phages have been shown to degrade the extracellular matrix of biofilms and gain access to underlying bacteria. This research explores the potential of bacteriophages to eradicate biofilms in an environment modeling a sink drain by comparing the efficacy, range, and durability of bacteriophage to a chemical disinfectant.

**Methods:** *E. coli* biofilms were grown in M9 minimal media placed in sink P-traps assigned one of three treatments: chemical disinfectant, bacteriophage, or deionized water (control). Biofilms were quantified at five time points -- 1, 12, 24, 36, and 48 hours -- using the crystal violet assay.

**Results:** Both chemical disinfectant and bacteriophage significantly decreased the optical densities of biofilms (p < 0.001***). P-traps treated with bacteriophages showed more uniform destruction of biofilm across P-trap compared to chemical disinfectant (p < 0.01**). A trend may suggest that over time bacteriophage became more effective at reducing biofilm than chemical disinfectant.

**Conclusion:** This work highlights the potential of bacteriophage as an alternative to conventional chemical disinfectants for biofilm control in settings such as hospital drains.

**Importance:** Nosocomial infections prolong hospital stay, costing the U.S. healthcare system $5-10 billion annually. An increasing number of reports demonstrate that sink drains -- reservoirs for multidrug resistant bacteria -- may be a source of hospital-related outbreaks. Recent studies have elucidated the mechanism of dispersal of bacteria from contaminated sinks to patients, but limited data are available identifying disinfecting methods for hospital drains. Not only did this study demonstrate that bacteriophages could reduce biofilms on sink drains just as effectively as a commercial disinfectant, it showed that phages tended to spread more thoroughly across P-traps and may work for longer. With hand-washing an imperative activity for disease prevention, hospital sinks should remain clean. This work explores an alternative disinfecting method for hospital sink drains.

## Introduction

In hospital settings, *E. coli* biofilms cause numerous medical-device-associated infections and recurrent urinary tract infections (Sharma et al., 2016). Biofilms have been shown to make *E. coli* less susceptible to antibiotics and biocides (Mittel et al., 2015). Several reports have found hospital sink drain pipes to be highly colonized with multidrug resistant bacteria (Kotay et al., 2017) (Lowe et al., 2012). One study found that cultures from handwashing sinks in the intensive care unit yielded *Klebsiella oxytoca*, an opportunistic pathogen that targets immunocompromised hospital patients (Lowe et al., 2012). Although intended only for hand hygiene, hospital sinks are also used for disposal of fluids, including body fluids, feeding supplements, and left-over beverages, which may act as nutrients to promote bacterial colonization and biofilm formation (Lowe et al., 2012). Moreover, the moist, closed environment of the drain promotes microbial growth (Kotay et al., 2017). In response to an outbreak at the National Institutes of Health Clinical Center, a study used green fluorescent protein expressing *E. coli* to reveal the mechanism in which bacteria from a sink drain reached hospital patients. They found that *E. coli* was able to form biofilms in a sink’s P-trap (fig. 1), the part of the sink drain that holds stagnant water to prevent the release of sewage gases (Kotay et al., 2017). The biofilms extended upwards to reach the sink strainer; once in the strainer, droplets containing *E. coli* can disperse to surrounding areas in the sink, reaching the hands of hospital patients and health care workers (Kotay et al., 2017).

**Figure 1.**
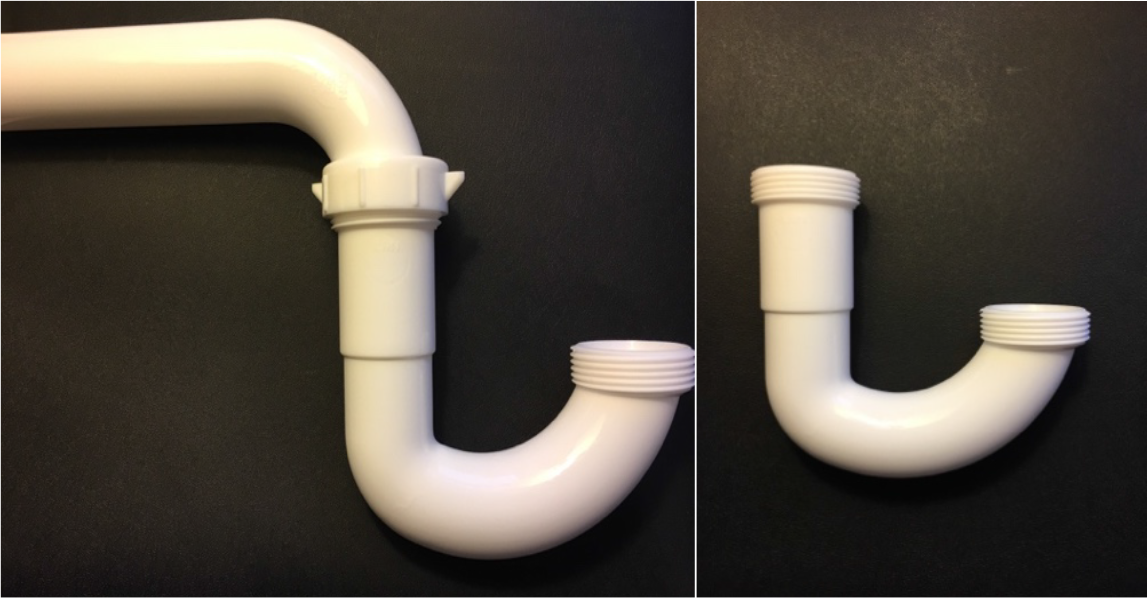
Side view of sink P-traps.

Bacteria present in biofilm states may show high levels of resistance to biocides and antibiotics (Harper et al., 2014). Furthermore, extensive use of chemical sanitizers may cause bacteria to develop resistance. In one study, the growth of *Acinetobacter baumannii* in ICU rooms was enhanced after the exposure to ethanol and alcohol-based cleaning products (Chen et al., 2013). Finally, although disinfectants are inexpensive, they are hazardous to the environment and contain compounds that may pose a danger to vulnerable patients (Ho et al., 2016). Common chemicals in disinfectants include sodium hypochlorite, a respiratory irritant and sense organ toxicant; phenols, which are recognized carcinogens; and ammonium compounds, which induce asthma attacks and contact dermatitis. Attempts at pouring strong chemical disinfectants -- such as bleach -- down sink drains have often been insufficient in killing bacteria (Kotay et al., 2017). Thus, there is a need to develop alternative disinfecting strategies in hospital environments.

While biofilms pose unique problems for antibiotics and biocides due to the extracellular matrix and the presence of metabolically inactive persister cells, bacteriophages, naturally occurring viruses that infect bacteria, have been found to prevent the formation of as well as reduction of mature biofilms (Harper et al., 2014) (Parasion et al., 2013). This is largely because phages have co-evolved with biofilm-forming bacteria (Harper et al., 2014). In biofilms, bacterial microcolonies surrounded by extracellular polymeric substances (EPS), the matrix and heterologous microbial cells may impede viral access to the bacterial cell surface (Sutherland et al., 2003). However, some phages are able to synthesize depolymerases that break down the EPS layer of the biofilm, allowing phages to enter the deeper biofilm layers to lyse susceptible bacterial cells (Parasion et al., 2014). Furthermore, many biofilms have open structures with water-filled channels that allow phage to enter the biofilm; one study demonstrated the radial movement of T4 phages across an *E. coli* biofilm, suggesting that the phages were not inhibited by any extracellular macromolecules present and may destroy biofilms in a single dose (Sutherland et al., 2004). Bacteriophages replicate within the host, releasing amplified numbers of progeny into the biofilm; as a result, phages can progressively spread through and remove the biofilm (Harper et al., 2014). Phages are also able to infect persister cells, remaining dormant within them, re-activating when bacteria become metabolically active (Harper et al., 2014). The large numbers of bacteria present within biofilms may even allow for more rapid and efficient phage amplification compared to planktonic systems. When 17 strains of planktonic *Pseudomonas aeruginosa* were tested to be insensitive to bacteriophages in a plaque assay, 8/17 strains supported the same bacteriophages when grown in biofilm states (Harper et al., 2014). Moreover, phages are highly specific to a particular bacterial species and are non-toxic to eukaryotic cells, making them a safe alternative to chemical disinfectants. While bacteria may grow resistant to chemical disinfectants, the selection process of phage-resistant bacteria was ten times slower than in the case of antibiotic-resistant bacteria (Parasion et al., 2014). Finally, phages increase in titer as they infect, which may allow for treatment to reach spaces in sink drains that are difficult to reach with conventional chemical disinfectants.

Phages have been used as environmental and consumer-friendly biological control agents (Le et al., 2017). In a recent study using *E. coli* and *Salmonella* phages against β-lactamase-resistant strains of *E. coli* on contaminated edible oysters, researchers found that phage treatments resulted in a significant decrease -- from 32.0×10^4^ CFU to 29.5×10^3^ CFU -- of *E. coli* (Le et al., 2017). Although phages have been widely studied as therapeutic agents in medicine and as biocontrol agents in food, limited data are present concerning the use of phages for disinfectant application and hospital hygiene. One study showed that bacteriophages protected medical devices such as catheters and prosthetic joints from colonization and biofilm formation (Fu et al., 2010). The pretreatment of catheters with phage reduced the bacteria to 4.03 log10 CFU cm^−2^ 24 hours after treatment (p < 0.001). Supplemental treatment with phage at 24 h also reduced biofilm regrowth (p < 0.001) (Fu et al., 2010). In a 2016 study, Ho et al. evaluated the potential of combining traditional cleaning practices with a bacteriophage containing aerosol. Phages targeting Carbapenem-resistant *Acinetobacter baumannii* (CRAB) were spread over the ICU room with an ultrasonic fogger containing phage suspension (Ho et al., 2016). The phages were able to spread to many hard to reach corners of the room. With a conventional alcohol and sodium hypochlorite-containing product alone, the specific strain of bacteria was still found to remain on hospital surfaces (0.6 CFU/cm^2^). Working alongside traditional disinfectants, phage containing aerosol successfully compensated for this deficiency. The mean percentage of drug-resistant isolates of CRAB also decreased from 87.76% to 46.07% after the introduction of phage (Ho et al., 2016). In another study working with phage ϕAB2, researchers showed that the addition of phage at a concentration of 10^7^ PFU/cm^2^ on a glass slide containing *A. baumannii* M3237 at 10^4^, 10^5^, or 10^6^ CFU/slide, significantly reduced bacterial numbers by 93%, 97%, and 99%, respectively (Chen et al., 2013). To show the versatile nature of the phage, they combined it with cream containing paraffin mineral oil to make a phage-based lotion to be used as hand sanitizer, which had the ability to reduce *A. baumannii* by 99% when spread over agar (Chen et al., 2013).

While limited data are available concerning the use of bacteriophages on hard surfaces, in lotions, and in soaps, phage application in hospital drains has not been studied. This study investigated the use of bacteriophages to combat biofilms in a sink drain through comparing the efficacy, range, and durability of bacteriophages to a commercial chemical disinfectant to combat biofilms grown in sink P-traps.

## Methods

### Experimental design

This study was conducted over five time points (Table 1), after the initial biofilm growth period (also 48 hours). The experimental unit is a sink P-trap. Time point zero marks the application of treatments: deionized water, chemical disinfectant, or bacteriophage. Additionally, a control group of P-traps were given only media with no *E. coli* inoculum to account for the background staining of crystal violet. At each of the time points shown below (Table 1) a P-traps were stained in order to quantify biofilm biomass after treatment. The sample size was 3 P-traps for each treatment per time point, making for a total of 45 P-traps. The sampling was destructive; each P-trap was disposed of after staining. For the P-traps in the 24 hour time group, photographs were taken of each side of the P-trap (side A, where the treatment was applied; side B, where no treatment was applied) in order to analyze the intensity of staining (pixels) using ImageJ. The response variables were the remaining biomass of biofilm after treatment (OD570 measurements) as well as average intensity in pixels to measure the spread of each treatment.

**Table 1.** Experimental design: the experiment was carried out over the course of 48 hours. Sampling was destructive; at each time point, a P-trap was taken and sampled. A total of 15 P-traps were tested for each of the three treatment groups (n=15).

|  | Traditional<br>Chemical<br>Disinfectant | Deionized water<br>(Control) | Phage-based<br>Disinfectant |
| --- | --- | --- | --- |
| 1 | n=3 |  |  |
| 12 |  |  |  |
| 24 |  |  |  |
| 36 |  |  |  |
| 48 |  |  |  |

### Phage and Bacterial Host Strain

Bacteriophage T4 was obtained from Carolina Biological in a stock solution with concentration of 2.5*10^10^ PFU/mL. *E. coli* K12 was obtained from Carolina Biological. Polyvinyl Chloride (PVC) Sink P-traps (model #25023, Keeney, CT) were obtained from Lowes.

### Solution Preparation

The traditional chemical disinfectant that was used was Clorox disinfectant (Item # 03050939, Clorox, CA) with sodium hypochlorite, sodium chloride, and sodium carbonate. To prepare the M9 minimal media, M9 salts were made using 64g Na_2_HPO_4_−7H_2_O, 15g KH_2_PO_4_, 2.5g NaCl, and 5.0g NH_4_Cl adjusted to 1000 mL with deionized water. For every liter of M9 media 200 mL of M9 salts were added to 800 mL of deionized water. To supplement the media, 5 mL of 20% glucose and 0.5mL of 1M MgSO_4_ were added. To make SM buffer, 10mL of 1M Tris pH 7.5, 10mL of 1M MgSO_4_, and 4g NaCl were added to 980 mL dd H_2_O. All solutions were autoclaved before use.

### Growth of Biofilms

To grow biofilms in sink P-traps, a culture of *E. coli* K12 was grown over night in LB media in a shaking water bath at 37°C. The overnight culture was diluted 1:100 into M9 minimal media supplemented with glucose and magnesium sulfate; this low nutrient medium slowed down the growth rate of cells in order to preserve the biofilm phenotype for the longest possible time. Seventy-five mL of the dilution were added into each P-trap. Parafilm covered both ends of the P-trap to prevent evaporation. To initiate biofilm growth, the P-traps were incubated for 48 hours at 37°C (fig. 2). A randomization process was done to assign each P-trap to a spot in the incubator.

**Figure 2.**
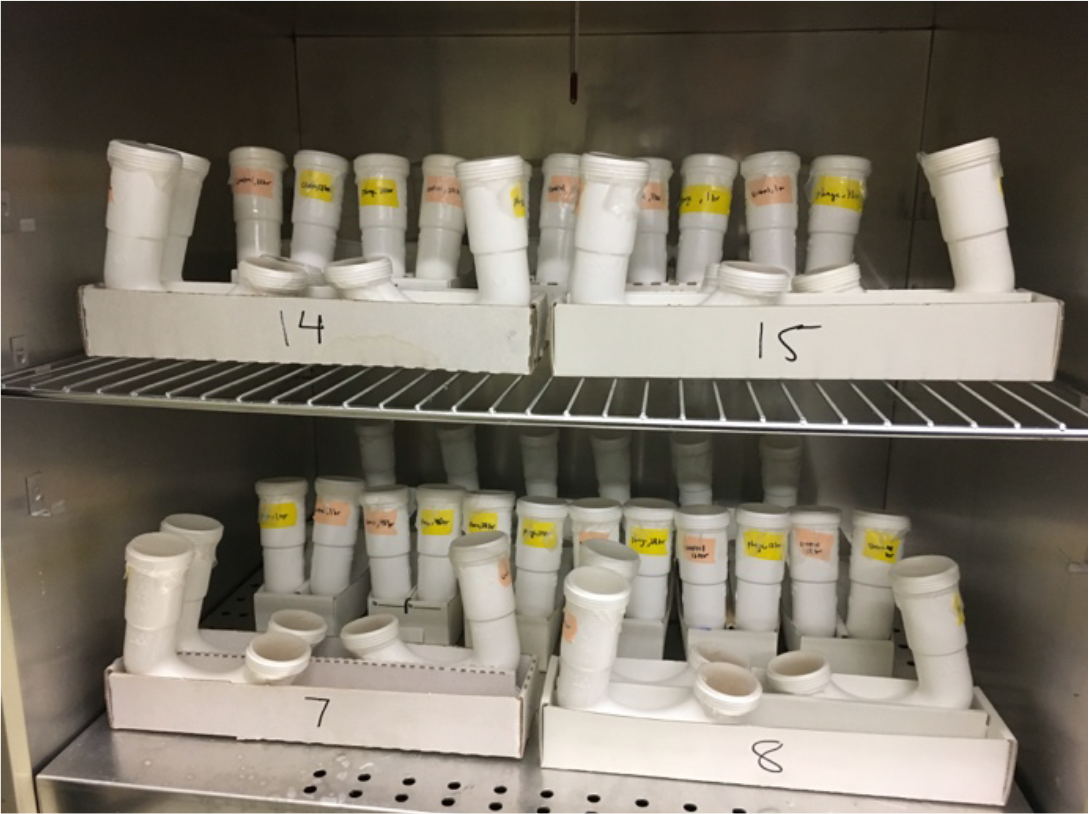
P-traps placed in an incubator for biofilm growth.

### Treatment of P-traps

After the biofilms had grown, P-traps were moved out of the incubator and on to a room temperature bench top, maintaining randomized positions. After the initial 48 hour period of biofilm growth 178 uL of each treatment was pipetted into opening A of the sink trap. The phage titer was 5*10^7^ pfu/mL. After the treatments had been applied, the sink traps were kept at room temperature.

### Biofilm Quantification Assay

To ensure all media components and planktonic cells were washed out, each sink trap was thoroughly rinsed three times with deionized water. Seventy-five mL of 0.1% crystal violet were added to each P-trap to stain the biofilm. After 15 minutes, the crystal violet was rinsed out 4-6 times with deionized water, shaking out and blotting vigorously on a stack of paper towels to rid the P-trap of planktonic cells and excess dye and reduce background staining. The 24 hour P-traps were photographed when dry. To quantify the optical density, 30 mL of 30% acetic acid were added into each P-trap to solubilize all of the crystal violet stain inside the P-trap. The solubilized stain was placed into cuvettes, and absorbance was quantified with a spectrophotometer (Spectronic 200) at wavelength 570 nm, using 30% acetic acid as a blank.

### ImageJ Photograph Quantification

Phages increase in titer as they infect, which may allow them to spread throughout the sink P-trap. In order to measure the ability of the treatments to spread, pictures were taken at both ends of the P-trap: at one end where treatments were applied (side A) and at the opposite end where no treatment was applied (side B). By using ImageJ, the brightness of the image was quantified using a mean intensity reading of an entire highlighted region – the opening of a P-trap (fig. 3). The smaller the mean intensity value, the less bright the highlighted area, which indicated greater amounts of biofilm. A difference between the pixel intensity of opening A and opening B were calculated to measure the ability for the phage to spread compared to the chemical disinfectant.

**Figure 3.**
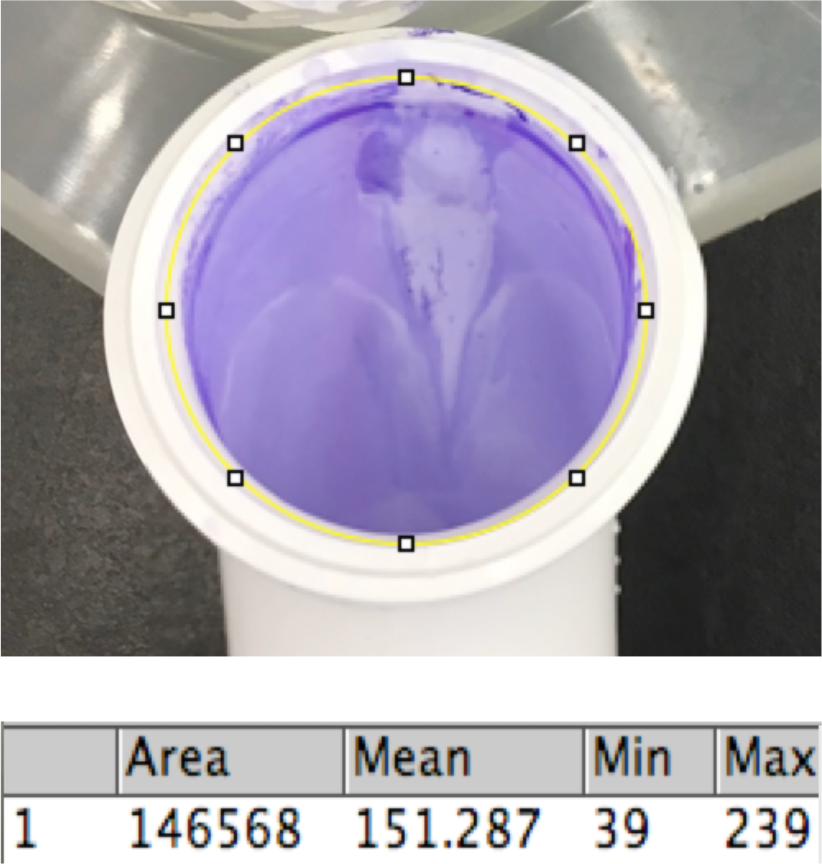
The average intensity in pixels (displayed as Mean) was quantified by fitting an ellipse using ImageJ; brightness correlates to the amount of crystal violet present, and thus the biomass of biofilm.

### Data Analysis

The two response variables in this study were OD570 (absorbance) and intensity in pixels (ImageJ). The statistical software JMP 10 was used for data analysis. Specifically, ANOVA and Tukey-Kramer Mean Separations were performed to analyze data.

## Results

### Growth of Biofilms

After a 48-hour incubation time period, biofilms grew successfully in the P-traps; this is evident through the presence of crystal violet staining via biofilm quantification assay in the deionized water treatment. Ring of heavy stain was observed near the air-liquid interface as expected as *E. coli* are motile. When comparing crystal violet stain of P-traps inoculated with *E. coli* to those which were not inoculated with *E. coli* (only M9 media), OD readings were significantly lower in P-traps containing no inoculum (fig. 4). Thus, the background staining of crystal violet on P-traps was accounted for and determined to be minimal.

**Figure 4.**
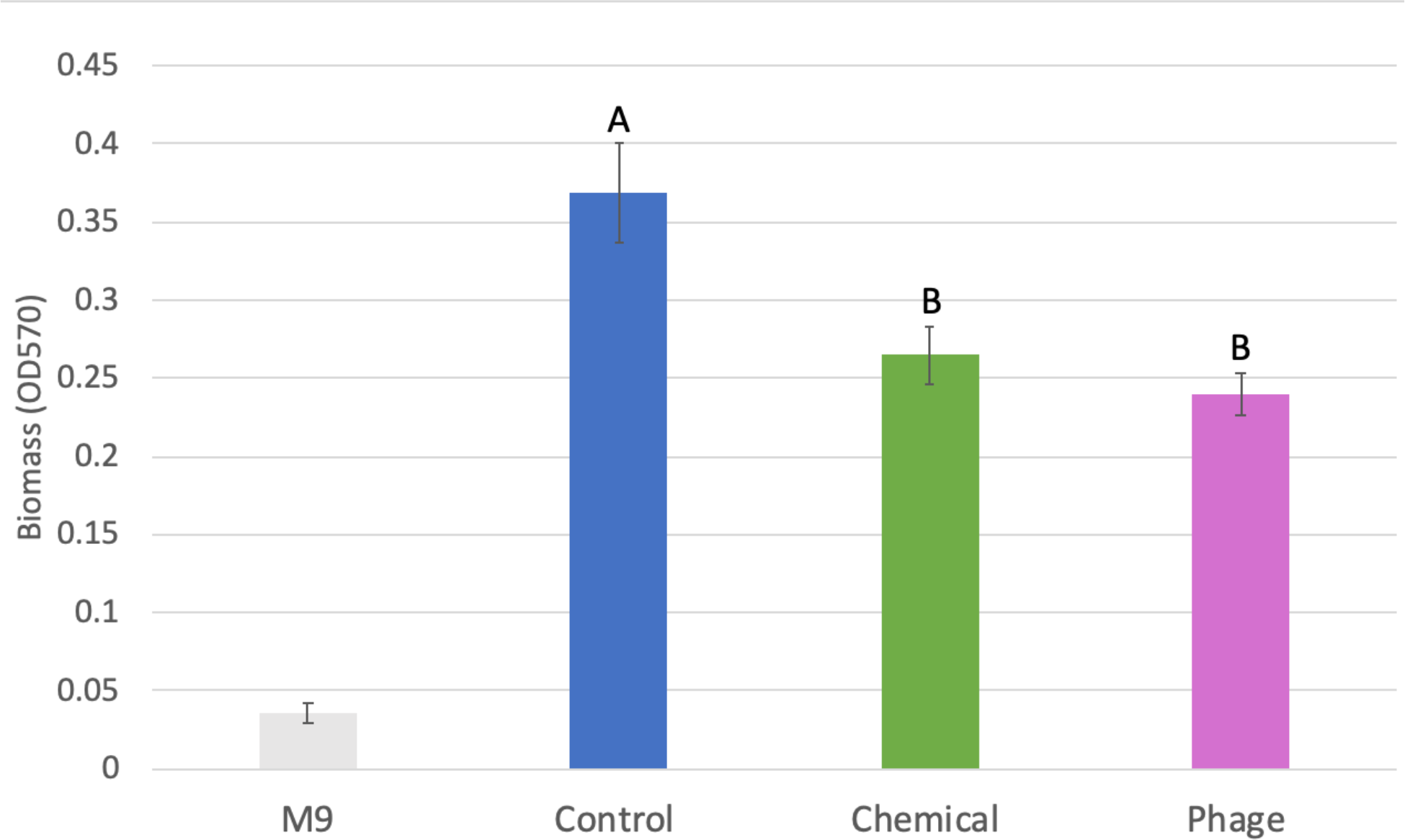
Biofilm Biomass after Treatment. The bar graph shows the average biomass of biofilms in P-traps containing M9 only (gray), treated with deionized water (blue), chemical (green), and phage (pink); each bar represents 15 P-traps taken at all times points. Both the chemical and phage were able to significantly decrease biofilm biomass (Table 2, ANOVA, df 2, p < 0.001***). One SE is shown. Letters indicate differences from Tukey Kramer mean separation.

### Biofilm Quantification Assay

P-traps treated with deionized water (control) showed significantly greater OD readings compared to P-traps treated with bacteriophage and chemical disinfectant (Table 2, ANOVA df 2, p < 0.001***). OD readings are proportional to the amount of crystal violet and biomass of the biofilm in the P-trap. As evident by these readings, both treatments were able to reduce biofilms (fig. 4). An ANOVA (Table 1) and LS Mean Differences Tukey-Kramer test (letters in fig. 4) were used to determine that this difference was statistically significant. The chemical disinfectant and the bacteriophage did not differ significantly from each other (fig. 4).

**Table 2.** An ANOVA of the optical density of biofilms over all time points. Significance is present between different treatment levels.

ANOVA
| Source |  | *df* | *SS* | *MS* | *F* | *P* |
| --- | --- | --- | --- | --- | --- | --- |
| **Model** |  | 5 | 0.16559245 | 0.033118 | 4.3675 | 0.003** |
|  | Time | 1 | 0.02305562 |  | 3.0405 | 0.0891 |
|  | Treatments | 2 | 0.13917604 |  | 9.177 | 0.0005*** |
|  | Time*Treatments | 2 | 0.00336079 |  | 0.2216 | 0.8022 |
| **Error** |  | 39 | 0.29573319 | 0.007583 |  |  |
| **Total** |  | 44 | 0.46132564 |  |  |  |

### Effectiveness of Treatment Over Time

While additional experiments should be conducted spanning longer time periods to detect a statistically significant effect of time, preliminary results suggest that bacteriophage may work more effectively for longer time periods than chemical disinfectant, as depicted in the trend of biofilm quantity over time (fig. 5). The biofilms in the control group remained consistently denser than the biofilms in the phage and chemical groups.

**Figure 5.**
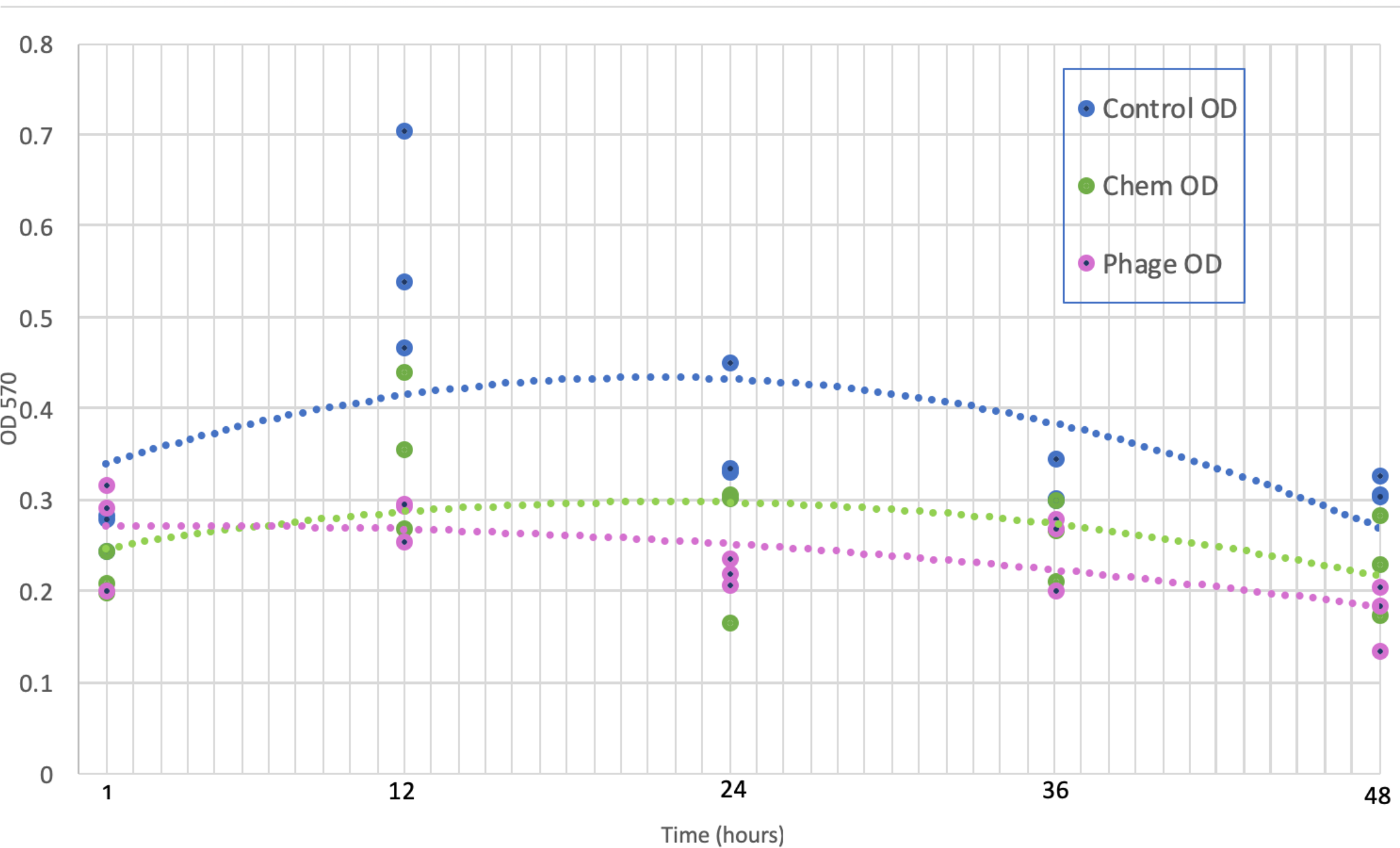
Effectiveness of Treatment Over Time. Scatter plot shows the trend of biofilm biomass at several time points after treatment. The blue shows the control (deionized water); the green shows the chemical disinfectant; and the pink shows the bacteriophage.

### Treatment at Early Stages of Biofilm Formation

An experiment was done to test the bacteriophage treatment at earlier stages of biofilm formation. One group of P-traps were exposed to phage immediately after bacterial inoculation, another group was exposed to phage when the biofilms were in the growth phase -- 90 minutes post bacterial inoculations (fig. 6). As expected, the P-traps treated with phage 0 minutes after bacterial inoculation showed significantly lower biofilm establishment (Table 3, ANOVA, df 2, p < 0.01**).

**Figure 6.**
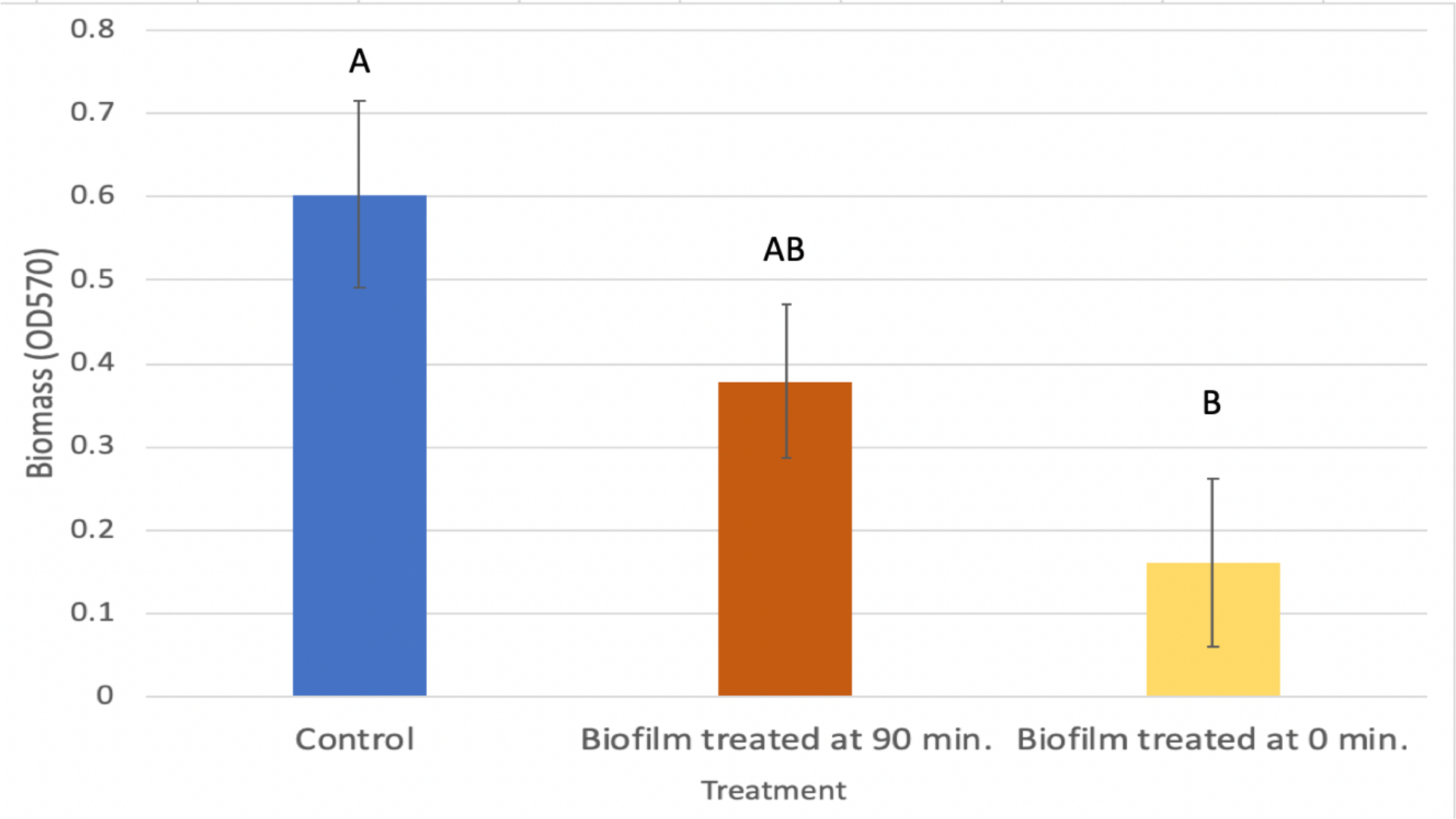
Treatment of Early Stage Biofilms. The bar graph shows the average mass of biofilms left in P-traps. One group of P-traps was given bacteriophage treatment 90 minutes post inoculation (orange) while another group was given the phage treatment 0 minutes post inoculation (yellow). One SE is shown. Lettering indicates differences from Tukey Kramer mean separation.

**Table 3.**
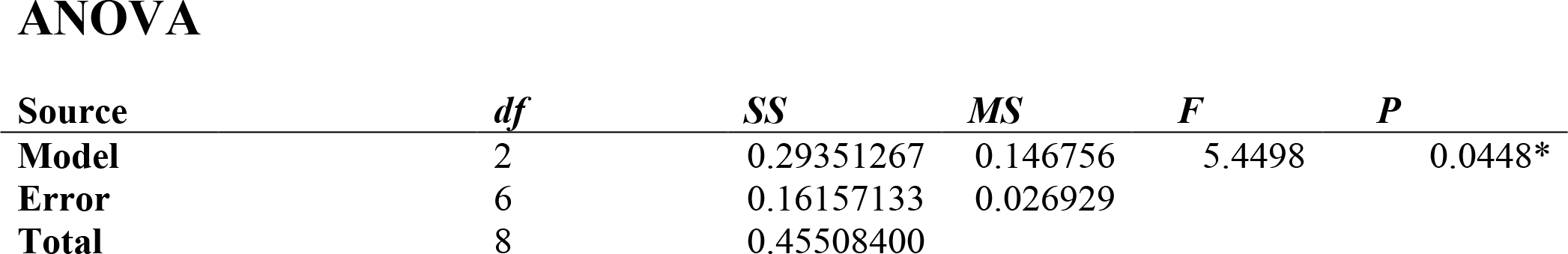
ANOVA test used to determine significance between the optical density of biofilms in P-traps treated at different stages of biofilm growth.

ANOVA
| Source |  | *df* | *SS* | *MS* | *F* | *P* |
| --- | --- | --- | --- | --- | --- | --- |
| **Model** |  | 2 | 0.29351267 | 0.146756 | 5.4498 | 0.0448* |
| **Error** |  | 6 | 0.16157133 | 0.026929 |  |  |
| **Total** |  | 8 | 0.45508400 |  |  |  |

### ImageJ Photograph Quantification

In order to measure the spread of the treatment in the sink P-trap, the brightness of both openings (opening A and opening B) of P-trap were measured using ImageJ (pixel intensity) 24 hours after treatment (fig. 7). The higher the pixel count in ImageJ, the denser the biofilm growth in that region (fig. 8). The intensity reading for the opening which the treatment was applied to was subtracted from the intensity reading of the opening opposite the side of treatment. The average difference (n=3) in intensity between side A and B for the control was 71.099 pixels (SE=3.15). The average difference in intensity for P-traps treated with the chemical was 76.588 pixels (SE = 2.41). The difference in intensity for P-traps treated with phage was the smallest, at 51.585 (SE = 4.52).

**Figure 7.**
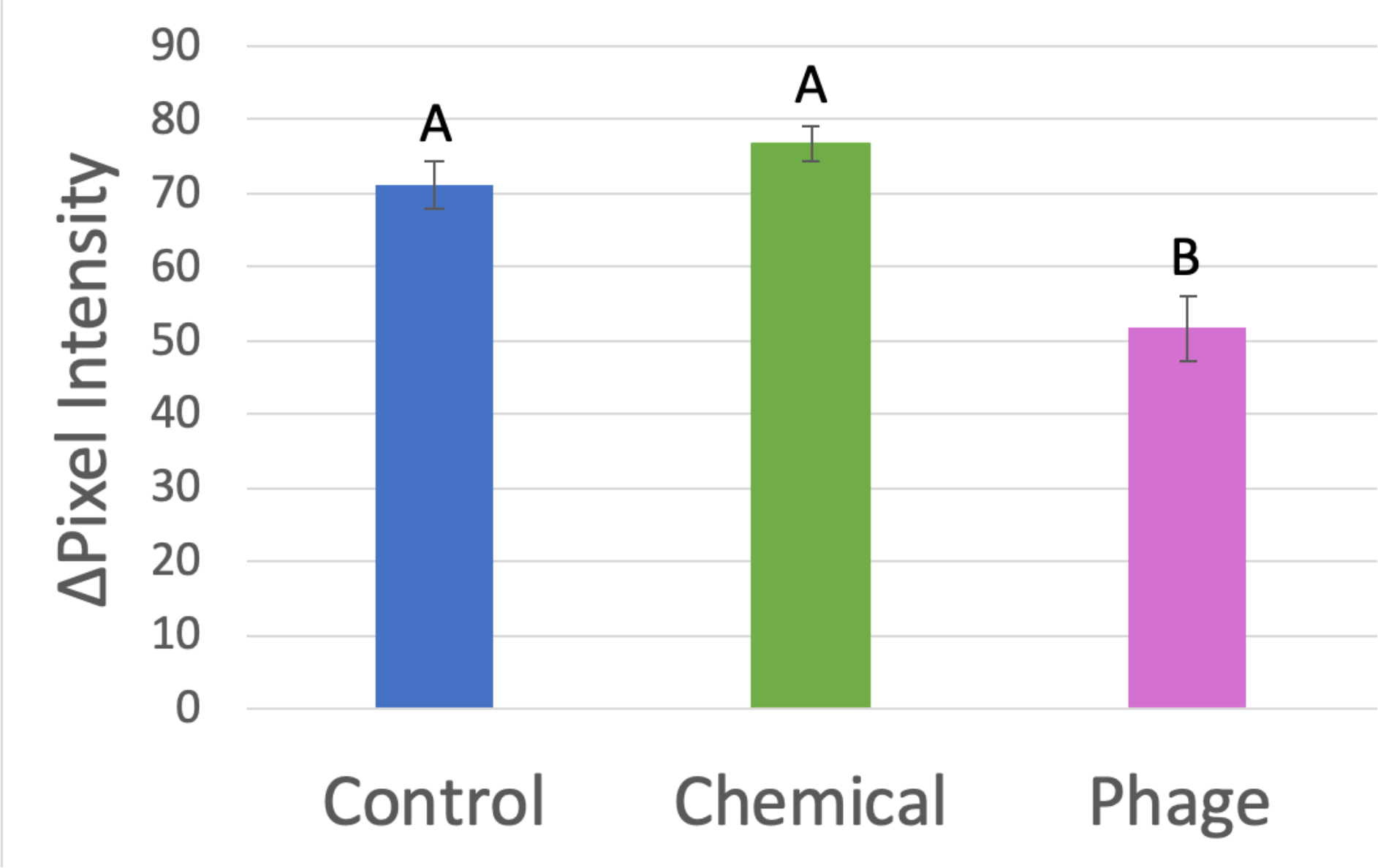
Spread of Treatment. This bar graph shows the difference in pixel intensity between the opening of the P-trap where treatment was applied and the opposite opening. The delta pixel intensity was significantly lower in the P-traps treated with bacteriophage (ANOVA df 2, p < 0.01**) (Table 4), suggesting that phage treatment was able to spread to a greater range throughout the P-trap compared to the chemical disinfectant. No significant difference was present between the control and chemical treatment group (letters indicate differences from Tukey Kramer mean separation.)

**Figure 8.**
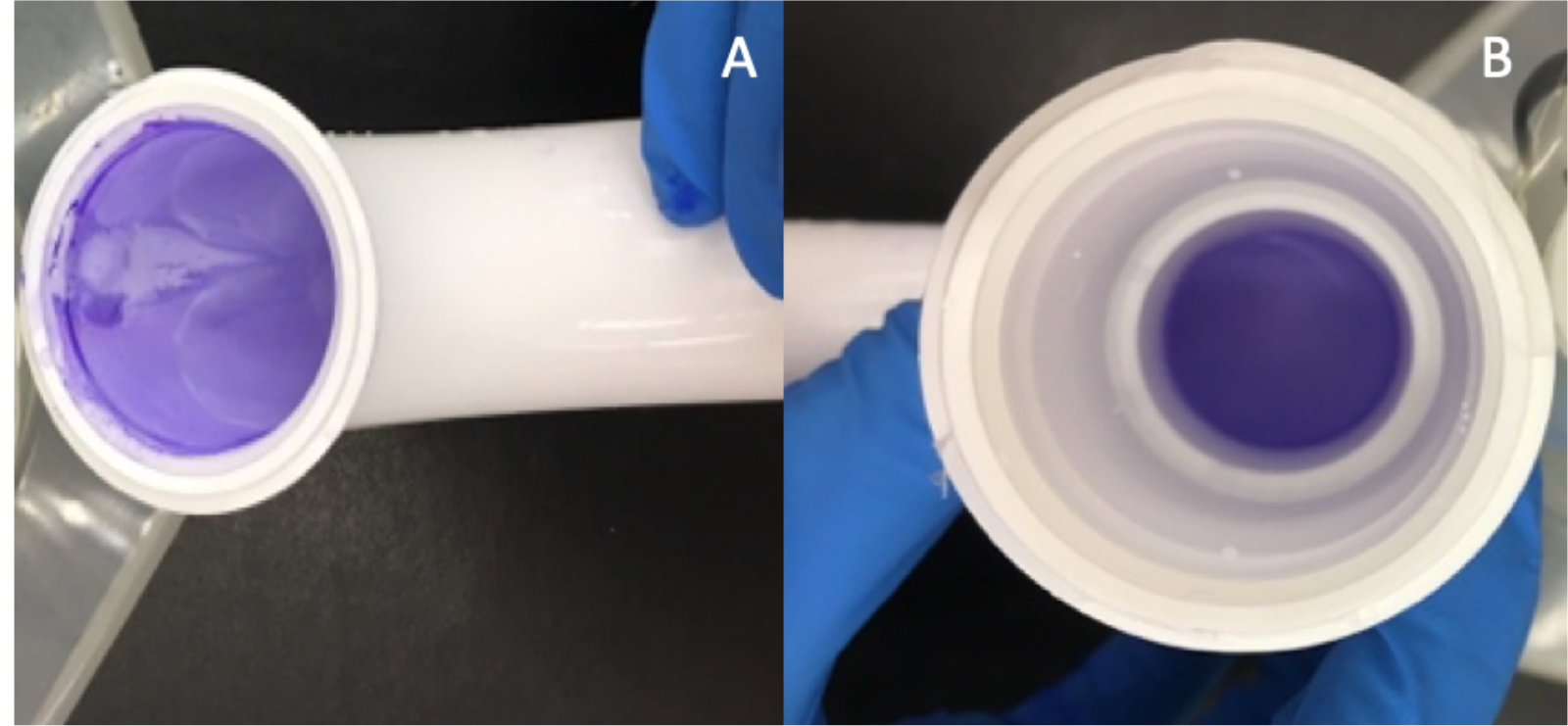
Photos of direct (Side A) and opposite sides (Side B) of a P-trap treated with chemical disinfectant. A visible streak appears on the side that the treatment was directly applied to, while the opposite side appears darker, indicating heavier biofilm growth.

## Discussion

The present study found that P-traps treated with bacteriophage as well as P-traps treated with chemical disinfectant showed a significantly lower OD570 reading compared to P-traps treated with a control of deionized water (Table 2, ANOVA, df 2, p < 0.001***). This indicated that both chemical disinfectant and bacteriophage T4 were able to reduce *E. coli* biofilms in Polyvinyl chloride sink P-traps and that phage was as effective as the chemical disinfectant.

Previous studies have tested phages in hospital-like setting, such as on the coating of a catheter to reduce the formation of a biofilm, but no previous study has tested bacteriophage on biofilms grown in a sink drain material. By applying bacteriophage directly to *E. coli* K-12 biofilms grown on PVC sink P-traps -- an environment intended to mimic a hospital sink drain -- this study builds on previous studies which show bacteriophage T4 can lyse *E. coli* K-12 biofilms. Furthermore, this study also compared the phage against a commercial chemical disinfecting product, Clorox bleach (NaOCl, Na_2_CO_3_, NaCl).

When looking at treatments across time, it was predicted that chemical disinfectant would be able to work in the short term, but that bacterial populations may start increasing shortly after the chemical disinfectant had evaporated. Making a prediction from the trend of biofilm OD taken in figure 5, bacteriophage disinfectant may remain effective for longer periods of time than chemical disinfectants; however, additional experiments should be conducted spanning longer time periods to detect a statistically significant effect of time. Biofilms in all three treatment groups began to decline after the 12-hour time point (fig. 5). A possible reason for the dwindling of biofilms in the control group over time may have been due to the lack of oxygen in the stagnant P-trap water. When the sink drains were stained 1 hour after treatment, no biomass reduction was observed in the bacteriophage treated sink traps. However, 12 hours after the phage challenge, a significant reduction of biomass was detected, indicating that the bacteriophage began producing noticeable effects between the 1 and 12-hour time points. The biofilm reducing effect happened sooner for P-traps which received chemical disinfectant, which showed a decrease in biomass at 1-hour post treatment. The treatments in this study were applied to mature biofilms that had been growing for 48 hours. Previous studies have demonstrated that bacteriophages lysed cells relatively slowly in biofilm states. The slower lytic performance of phages in biofilms is likely due to the physiological state of cells rather than the physical barrier the biofilm matrix (Cerca et al., 2007). The low metabolic activity of cells in mature biofilms are less susceptible to lysis while the high metabolic activity of bacteria in the initial attachment and early maturation stages of biofilm development are more sensitive to cell lysis (Cerca et al., 2007). Nevertheless, phages were able to lyse bacteria in mature *E. coli* biofilms.

Bacteriophages are replicating biological control agents, making them inherently different from chemical agents (Harper et al., 2014). This study hypothesized that bacteriophage would spread through a larger range of the P-trap than the chemical disinfectant because phages increase in titer as they infect (Ho et al., 2016). Using ImageJ pixel intensity measurements, it was found that the difference in intensity was significantly lower (Table 4, ANOVA, df 2, p < 0.01**) in the P-traps treated with bacteriophage than with chemical, which may indicate that the phage treatment was able to spread to the opposite end of the P-trap (fig. 6). However, in the control group, the difference in intensity was not significantly lower than the difference in intensity for the chemical treatment group. One explanation could be due to a small sample size (n=3), as there was one control P-trap that showed a particularly high difference in intensity, which brought the entire average up.

**Table 4.** ANOVA test used to determine significance between the differences in pixel intensities on ImageJ. The model indicates that the range of spread between treatments varied significantly.

ANOVA
| Source |  | *df* | *SS* | *MS* | *F* | *P* |
| --- | --- | --- | --- | --- | --- | --- |
| **Model** |  | 2 | 1036.0660 | 518.033 | 14.3437 | 0.0052** |
| **Error** |  | 6 | 215.6941 | 36.116 |  |  |
| **Total** |  | 8 | 1252.7600 |  |  |  |

This study showed that bacteriophages could be promising biocontrol agents to be used in hospital sink drains. Moreover, bacteriophages are relatively fast and inexpensive to manufacture (Parasion et al., 2014). Much work, however, still needs to be done to implement bacteriophage into a real hospital setting. As sink drains lead to sewer systems which lead to a larger environment, studies should be done to examine the downstream effects of releasing bacteriophages into the environment. One limitation of this study was the lack of assessment for phage resistance. A concern with therapeutic uses of phages is the possibility that persister cells present in biofilms – dormant cells that neither grow or die in the presence of antimicrobials – could develop phage resistance (Hanlon, 2007) (Fu et al., 2010). Bacteriophages are diverse organisms, outnumbering bacteria in almost all environments and making up the most dominant life form in the biosphere (Parasion et al., 2014). This means that phage cocktails may be created to contain a diverse mixture of bacteriophages in order to slow the process of phage resistance up to ten times slower than antibiotic resistance (Parasion et al., 2014). A benefit of creating diverse phage cocktails would be to produce a disinfectant capable of targeting a wide range of host bacteria, a solution suitable to the hospital drain environment where multispecies biofilms tend to form (Fu et al., 2010) (Sutherland et al., 2004). It has also been shown that intervention with phage treatments may decrease the number of drug-resistant isolates of *A. baumanii*, a benefit that comes with using phage over chemical disinfectants in hospitals (Ho et al., 2016).

In conclusion, this was the first study to use bacteriophage to reduce *E. coli* biofilms in sink drains. Bacteriophage was found to reduce the mass of the biofilm and to spread to a greater range in the pipe than a chemical disinfectant.

## Acknowledgements

I would like to acknowledge Dr. Amy Sheck for assistance in the project development and for being a mentor; Dr. Kim Monahan for guidance during the Glaxo Summer Research Fellowship; Mr. Philip Rash for assistance in data analysis; the Glaxo Endowment to the North Carolina School of Science and Mathematics for funding the project; Research in Biology mentors: Madeline Paoletti, Elizabeth Farmer, Tony Zhang, Isaac Poarch, Billy Ngo, Tyler Edwards; Colleagues and Glaxo fellows Aarushi Patil, Abigail Philips, Emile Charles, Isabel Huesa, and Marco Allen.

